# A general framework explaining variation in plant economics traits with environment and through ontogeny

**DOI:** 10.64898/2026.05.25.727577

**Authors:** Daniel S. Falster, Isaac R. Towers, Peter A. Vesk, Mark Westoby

## Abstract

Plant economics traits, such as leaf mass per unit leaf area (LMA) and stem specific density (SSD), capture diversity among plant species in how common tissues (leaf, wood, root) are constructed. These traits are key descriptors of plant strategy, yet it has proven difficult to explain this variation with theory and process-based models. Here we reveal a general explanation on why these economics traits vary with environment, through ontogeny, and with other plant traits. This explanation relies on three core assumptions: 1) plants seek to maximise growth rate, 2) growth rate can be decomposed into a product, and 3) there is a tradeoff between the efficiency of tissue construction and tissue turnover rate. Formulation of growth as a product is essential, as it causes the optimal value of an economics trait to vary with the plant’s biomass production rate, which means economics traits will naturally covary with the abiotic environment, the competitive context, and other strategical features of the plant. Finally, we show how a modification of the trait into plastic and non-plastic components alters the magnitude of intra-specific responses, aligning model responses with empirical trends. Broadly, our results help explain how plant form and function for a wide diversity of species is shaped to suit their environment and, moreover, they reveal insight into a general fast-slow spectrum (Reich 2014) with coordinated shifts among organs (leaf & stem) through tradeoffs between efficient tissue construction and turnover.

## Introduction

Plant economics traits, such as leaf mass per unit leaf area (LMA) and stem specific density (SSD), capture diversity among plant species in how common tissues (leaf, wood, root) are constructed. A unit of leaf, wood or root built to be more robust inevitably requires more investment from the plant, leading to a trade-off between construction cost and turnover rate of parts. During recent decades, the economics traits LMA and SSD have emerged as some of the most important descriptors of plant strategy (1; 2; 3; 4), varying at multiple scales–including among coexisting species, across resource gradients, and through the ontogeny of individuals (Table 1). While such variation is thought to reflect adaptive responses, and is what makes the traits useful indicators of plant strategy and function, explaining why these traits vary simultaneously at multiple scales has remained a challenge.

**Table 1:**
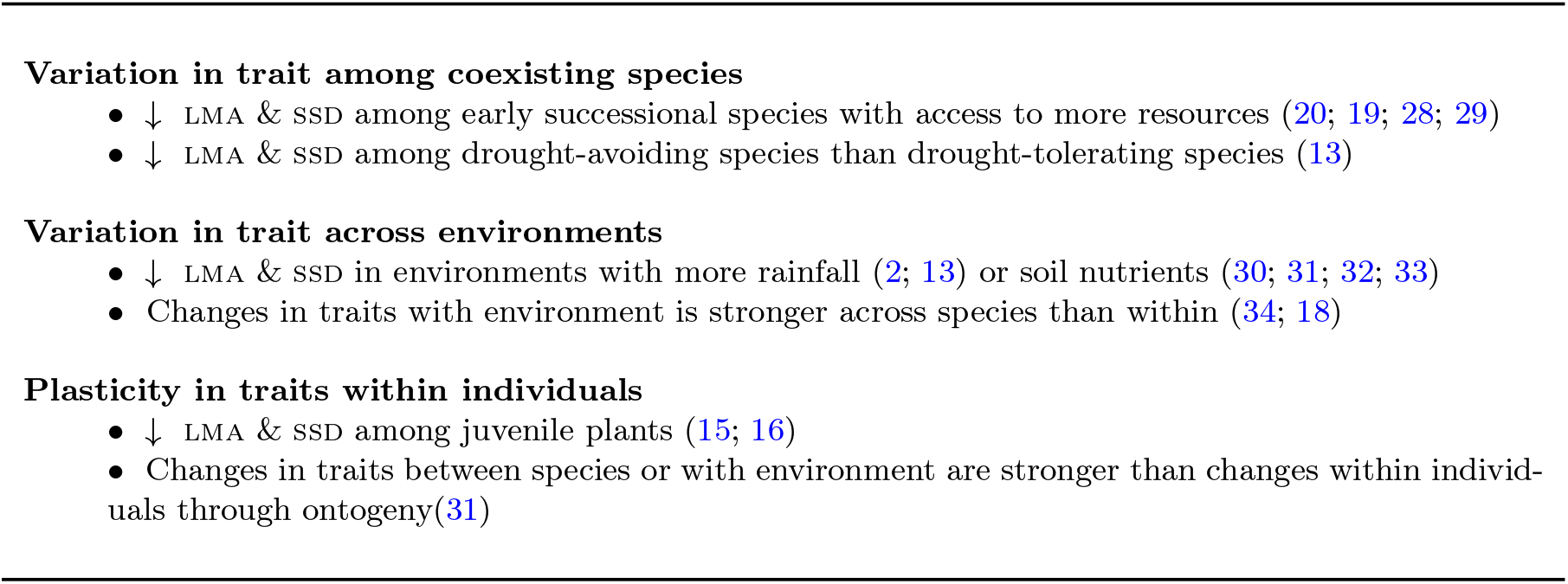
Observed phenomena predicted by theory optimising economics traits for plant growth rate.

Eco-evolutionary optimisation models (EEO) seek to explain plant responses to environment (5), by asking what values along a trait-based trade-off optimise fitness, and how this trait value changes depending on context. EEO models focusing on the net carbon budget of plants are most common (6; 5; 7, e.g.), and have succeeded in explaining multiple responses of leaves to environment. However, variation in economics traits has proven more difficult to explain. Indeed, in the absence of a direct emergent effect from the model, a recent analysis resorted to making phenomenological links between economics traits and environment (8). Here, we show how shifting the optimisation criteria of the EEO model from biomass production to the growth rate of a plant leads to a generic explanation for the response of economics traits to diverse contexts (Table 1), including variation through ontogeny, variation among coexisting species, and variation across environmental gradients. Moreover, we show that these responses emerge because the net effect of economics traits on growth rate operates via a product of two opposing effects, and products enable richer feedbacks in optimisation problems than do functions based on a difference, like biomass production.

A crucial advance in our analysis is the formulation of an individual’s growth rate as a product of two terms, together capturing a tradeoff between two opposing effects of an economics trait on plant growth: efficient construction vs increased turnover of parts. The standard approach for an EEO is to find the trait value maximising some fitness function. For example, for trait *x* and fitness *f* (*x, H, E*), dependent on *x*, an environmental variable *E* (e.g. amount of light, water, nutrients, or temperature), and individual’s ontogeny, here represented by height *H*; find the optimal trait value *x* = *x*^∗^ by solving 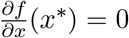. The science lies in identifying a biologically meaningful *f*, such that *x*^∗^ responds to other variables like *E* and *H*. Biomass production rate is often expressed as the difference between gains (photosynthesis) and loss (respiration, turnover), which implies the algebraic form for *f* is a difference (*f* (*x, H, E*) = *g*(*x, H, E*) − *h*(*x, H, E*)) and hence *x*^∗^ occurs when

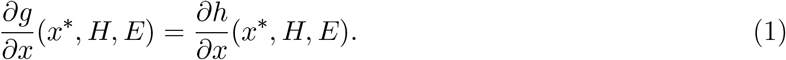

However, two barriers prevent general models for economics traits arising when *f* is a difference. First, there is no evidence that both the benefits and costs of economics trait can be expressed in terms of biomass production. SSD, for example, does not logically have effects on the benefit component of biomass production (photosynthesis). Second, only the derivatives of *g* and *h* inform *x*^∗^ (via Eq. 1), meaning any general additive effects of *E* or *H* on *g* or *h* don’t influence *x*^∗^.

By contrast, if *f* is a product, i.e. *f* (*x, H, E*) = *g*(*x, H, E*) × *h*(*x, H, E*), the values of *g* and *h* themselves become part of the solution, as the optimum *x*^∗^ must satisfy

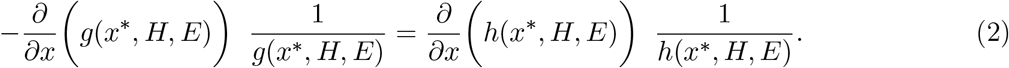

Eq. 2 includes *g* and *h*, and their derivatives, whereas only the derivatives are in Eq. 1. Thus, any factor influencing *g* or *h* could influence *x*^∗^. This result has far reaching consequences. As we show below, this feature of a fitness product enables any factor moderating individual biomass production or plant construction cost (like rainfall, height, or other traits) to influence the optimal value of an economics trait.

To apply this insight to economics traits, we leverage a model for plant growth rate from Trait-Growth-Theory (TGT), which decomposes growth rate (in leaf area *A*_*l*_ per unit time) for a plant with economics trait *x*_*i*_ (where *i* indicates leaf, bark, sapwood or roots), as a product of two terms (9; 10):

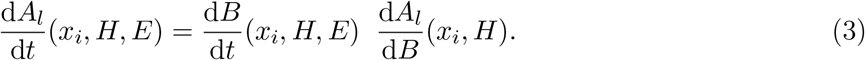

(See Table 2 for variable definitions.) The first term on the right is the net biomass production of the plant, which is assumed to vary with *x*_*i*_, *H* and *E*. The second term 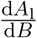 is the efficiency of leaf area deployment, which accounts for the marginal cost of deploying an additional unit of leaf area, including construction of the leaf itself and supporting bark, sapwood, and roots. The inverse of this term, 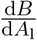 is the whole-plant construction cost per unit leaf area, which can be further decomposed as a sum of construction cost per unit leaf area for different tissues (9; 10) to yield:

**Table 2:** Variable definitions. All variables apply to an individual plant.

| Symbol | Unit | Description |
| --- | --- | --- |
| **Economics traits** |  |  |
| $x_1$ | $\text{kg m}^{-2}$ | Leaf mass per area (LMA) |
| $x_2$ | $\text{kg m}^{-3}$ | Wood density (SSD) |
| **Size metrics** |  |  |
| $H$ | m | Height of a plant |
| $M_i$ | kg | Mass of tissue type $i$ retained on plant |
| $A_l$ | $\text{m}^2$ | Area of leaf |
| $\gamma_i$ | $\text{m}^{2(3)} \text{ m}^{-2}$ | Area (or volume) of tissue $i$ per unit leaf area |
| **Environmental variables** |  |  |
| $E$ | | Resources available |
| **Rates of change** |  |  |
| $dA_l/dt$ | $\text{m}^2 \text{ yr}^{-1}$ | Growth rate of leaf area |
| $dB/dt$ | $\text{kg yr}^{-1}$ | Net biomass production |
| $dA_l/dB$ | $\text{m}^2 \text{ kg}^{-1}$ | Efficiency of leaf area deployment |
| $dB/dA_l$ | $\text{kg m}^{-2}$ | Whole-plant construction cost per unit leaf area |
| $\frac{\partial}{\partial x_i} (dB/dt)$ | $\text{m}^{2(3)} \text{ yr}^{-1}$ | Change in Net biomass production per unit increase in trait |
| $\frac{\partial}{\partial x_i} (dA/dB)$ | $\text{m}^{4(5)} \text{ kg}^{-2}$ | Change in efficiency of leaf area deployment per unit increase in trait |
| $P$ | $\text{mol yr}^{-1} \text{ m}^{-2}$ | Photosynthetic rate per unit area |
| $R_i$ | $\text{mol yr}^{-1} \text{ m}^{-2(3)}$ | Respiration rate per unit area (or volume) of tissue $i$ |
| $k_i$ | $\text{yr}^{-1}$ | Turnover rate for tissue type $i$ |
| **Other parameters (constant throughout)** |  |  |
| $\alpha_y$ | | Yield, fraction of carbon fixed into biomass |
| $\alpha_{\text{bio}}$ | $\text{kg mol}^{-1}$ | Biomass per mol carbon |

For $M_i$ , $R_i$ , $k_i$ and $A_i$ , subscripts refer to: l = leaves, b = bark, s = sapwood, h = heartwood, r = roots, st = total stem, a = alive tissue. For units with a bracketed exponent, these refer to the equivalent units for SSD, as opposed to LMA.

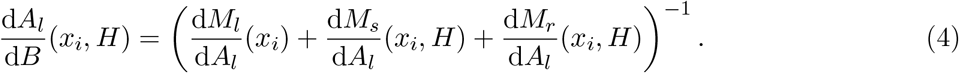

Eq. 4 captures the entire cost of building a unit of leaf. It is also where the benefit of deploying tissue cheaply, through changes in economics traits, is naturally expressed. The leaf component, (d*M*_l_*/*d*A*_l_), is the economics trait LMA, while the stem component increases with SSD, *H* and Huber value (the area of sapwood per leaf area). Allocation to roots also increases with *A*_*l*_. The model of biomass allocation captured by Eq. 4 differs from the standard approach in many vegetation models, where biomass allocation is modelled as a fixed fraction independent of economics traits (11). By contrast, in Eq. 4 building cheaper tissues allows plants to spend less on that tissue, with the savings gains reinvested in growth of the entire plant. This assumption is supported by a large scale empirical analysis showing that having a lower LMA leads to less leaf mass but no less leaf area (12); the opposite of what the standard models assume.

As Eq. 3 is a product, the optimal value 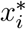 of economics trait satisfies Eq. 2, i.e.:

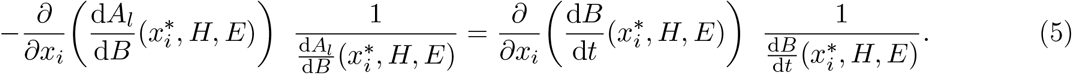

Eq. 5 has a natural interpretation; considering changes that arise from building a more expensive tissue, 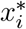 occurs where the relative decrease in the efficiency of leaf area deployment equals the relative increase in biomass production. Further, by optimising growth rate we also capture results from any previous model focussing on biomass production, as a special case. (If the trait in question only affects biomass production and not construction, the 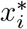 optimising biomass production also optimises growth rate.)

Crucially, the fact that Eq. 5 includes 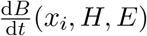 means that *x*^∗^ will shift with any factor that alters the net biomass production of the plant. This includes other plant traits, such as leaf-level photosynthesis, and environment, such as levels of light, water, or nutrients. The optimum trait shifts because the marginal value of building cheaper tissues increases with the overall productivity of the plant. Similarly, Eq. 5 causes the optimal trait to vary with any factor influencing 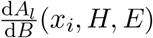, such as *H*. The fact that optimal economics traits respond to other traits, environment and size, without having any direct effect on the trait-based tradeoff itself, potentially drives responses of both LMA and SSD to a wide range of conditions, including with environment, among coexisting species, and through ontogeny (Table 1).

This result explains why LMA tends to decrease across sites as rainfall or temperature increases (13; 2), as such environments broadly allow for greater maximum productivity per plant. Fig. 1 shows that the optimal LMA decreases as the plant’s biomass production increases, and how the terms in Eq. 5 vary with the average photosynthesis per leaf area, 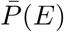. Changes in 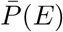 could reflect either varying resources (light, water) or variation in other traits within the plant (E.g., maximum photosynthetic capacity or hydraulic conductivity). Note how only the far-right-panel (biomass production) is varying with 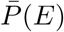, yet this is sufficient to shift the optimum. The optimum LMA is lower when 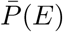 is high, because a high 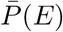 diminishes the potential costs of decreasing LMA, relative to the benefits. Thus species specialising on high resource environments within or across sites tend to have low LMA (Table 1). Such species may also have high potential photosynthesis or large vessels, which would further selection for low LMA (13).

**Figure 1:**
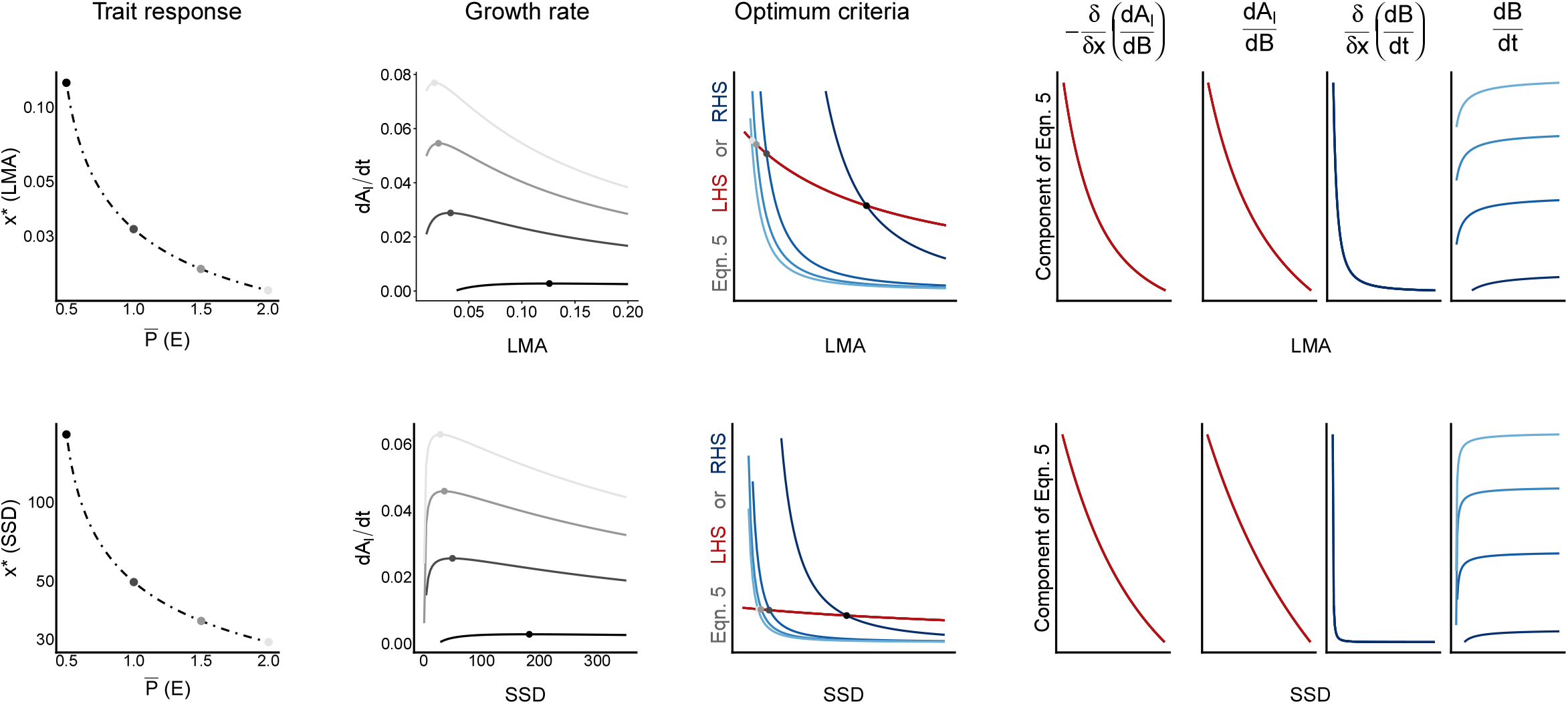
Optimal response of economics traits to changes in plant productivity. Rows show results for LMA (top) and SSD (bottom) respectively. First column shows change in trait optima with rate of individual photosynthesis 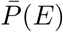. The points represent trait optima at four samples along the environmental gradient which are used for illustrative purposes in the following columns, with lighter shading indicating more productive environments. Remaining columns show changes of various outcomes with trait value, for individuals experiencing a given 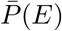 (as indicated by colour and corresponding to location of points in the first column). Second column shows growth rate (Eq. 3), with optimum represented by points. Third column shows either side of optimality criterion in Eq. 5, with *x*^∗^ occurring where the relative decrease in the efficiency of leaf area deployment (in red, the LHS of Eq. 5) intersects with the relative increase in biomass production (in blue, the RHS of Eq. 5). Columns 4-7 show the four elements in Eq. 5. (See main text for interpretation.) Note that variation in the position of *x*^∗^ with productivity emerges exclusively from upward shifts in net biomass production (column 7), 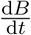, as other columns (4-6)don’t vary with 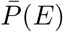.

**Figure 2:**
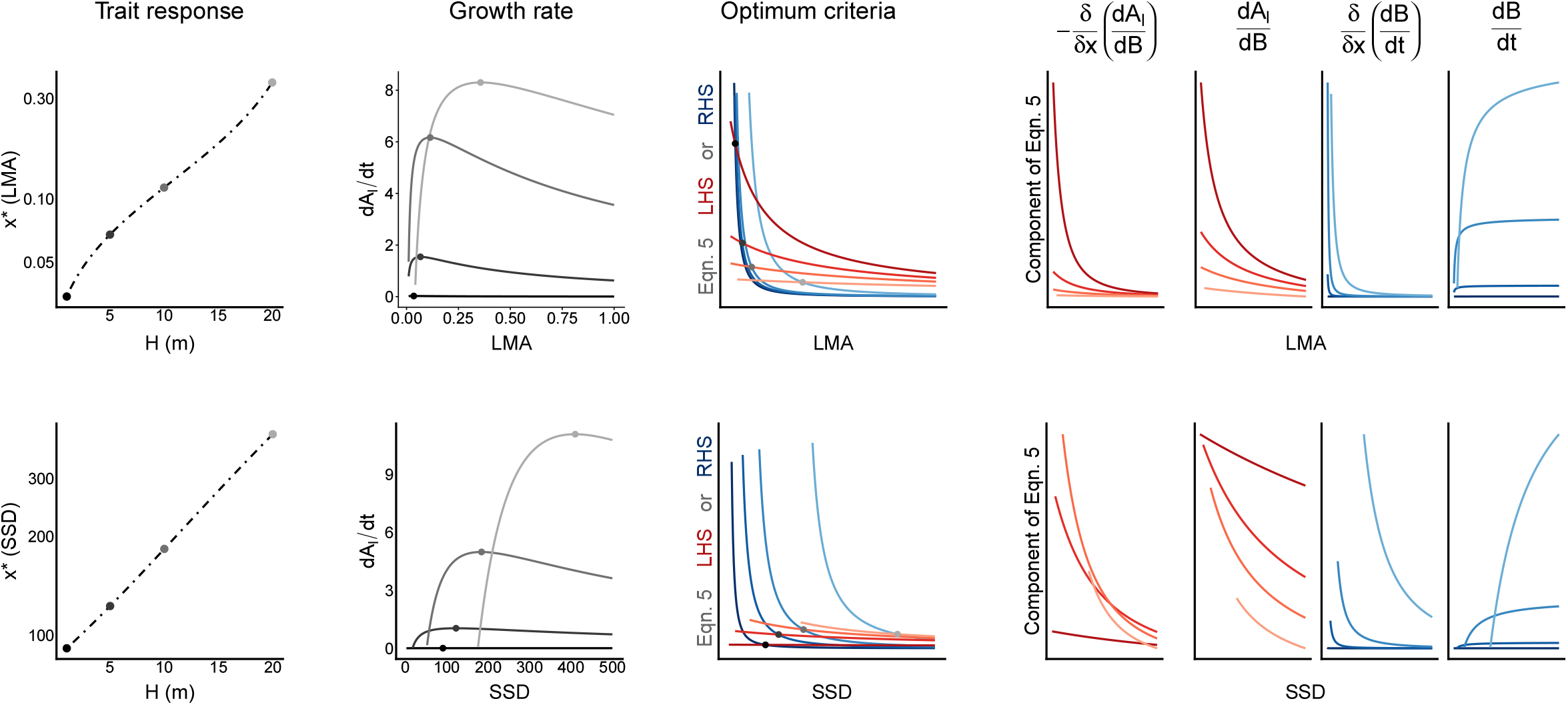
Optimal response of economics traits to changes in plant size. Panels and columns are identical to Fig. 1, except that in this figure different points and lines correspond to individuals with different *H* (lighter colouring indicates greater *H*), whereas in Fig. 1 they differed in 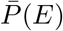. Note that varying *H* influences each term of Eq. 5.

Broadly, our analysis predicts lower SSD will be selected under similar circumstances to low LMA (Fig.1). Like LMA, lower SSD incurs a cost on biomass production, due to higher turnover, but will still be selected for when 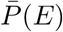 is high, as it facilitates rapid growth by increasing leaf deployment efficiency. It is common for fast growing pioneers growing under high light or post disturbance to have light wood (Table 1). SSD also increases in drier environments, among trees and shrubs (13). In arid areas, trees and shrubs must endure and grow during drier periods, and this is achieved by minimising turnover of stem through high SSD. In the same environments, other species show low SSD, however these drought avoiders grow during wet periods when resources are abundant (high 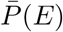 and retreat to the seed bank as productivity drops. Combined, the selection pressures for LMA and SSD suggest some coordination between leaf and stem economics traits. Such broad scale coordination has been observed, both in coordinated shifts of traits across climates (13), and in large global databases (4). Looking at correlation of traits worldwide, LMA and SSD correlate and form a major axis of variation (4).

Our framework also provides a general explanation for covariation between economic traits beyond independent responses of both traits to the environment. The EEO model predicted that, in a fixed environment, individuals with larger SSD have should have a larger LMA (Fig. 3). The reason that is occurred is because, at higher values of SSD, the cost of a unit increase in LMA to growth rate in terms of 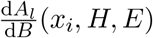 is comparatively lower relative to the benefit received in terms of greater 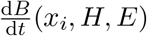 incentivizing individuals to invest in larger LMA. In simpler terms, if selection causes one economic trait to evolve towards more conservative strategies, other economic traits should also experience selection towards conservative strategies because it is inefficient to have a combination of fast and slow traits, thereby supporting the hypothesis that taxa converge on a uniformly fast, medium or slow strategy for all organs (14).

**Figure 3:**
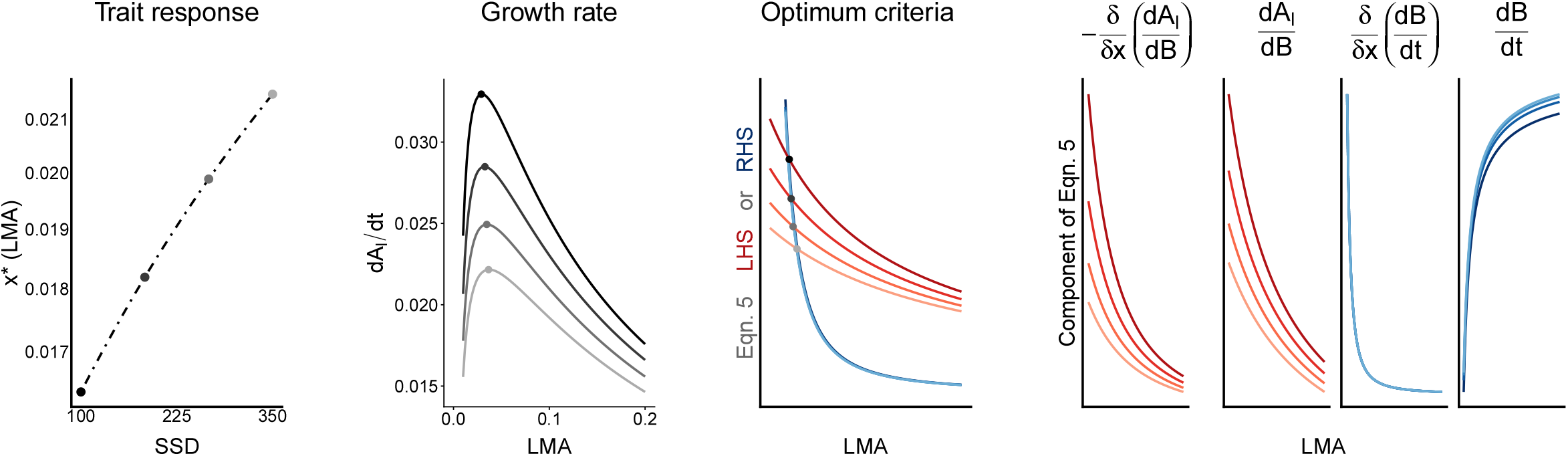
Optimal response of lma to changes in ssd. Panels and columns are identical to Fig. 1, except that in this figure different points and lines correspond to individuals with different SSD (lighter colouring indicates greater SSD), whereas in Fig. 1 they differed in 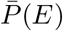. Note that varying SSD influences each term of Eq. 5.

The same EEO model predicts changes in traits through the lifespan of an individual, as a taller *H* modulates the relative costs and benefits of economics traits. Changes with *H* are more complex than changes with 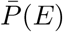, as changes in *H* change all four terms in Eq. 5 (Fig. 2). However, changes in *H* do result in simple predictions that broadly align with data (Table 1). Eq. 5 predicts that both LMA and SSD increase as plants increase in size (Fig. 2). The predicted changes in LMA are widely observed (15; 16). Likewise, we predict that SSD should increase, purely based on economics (Fig. 2) and without considering any implications for support. Indeed, fast growing species commonly increase in SSD as they grow large (17). By contrast, slow-growing shade -tolerant rainforest species may decrease in density as they grow larger (17). Superficially, this result goes against our prediction, except that, in such species, changes in height are confounded with changes in light. As saplings, shade tolerants may spend decades in low light conditions, where we would predict a high density to minimise turnover. They only begin to grow once a gap opens up. The increased light available would then decrease the optimal SSD, as observed.

A counter-intuitive feature of the results presented thus far is that they would suggest responses within species could be equal in magnitude to those observed across species. However, empirical data suggest weaker responses. For example, global variation in SSD responses to aridity suggest responses within species are 70-80% weaker than across species (18). Similarly, LMA will commonly double through ontogeny as an individual grows (15), but increase 10 times across species with aridity (13), or vary five-fold among coexisting species utilising different light environments (19; 20).

One plausible mechanism that could moderate the within-species response of traits would be if there were a cost to plasticity (21). When an additional cost of plasticity is imposed, on top of the underlying biophysical tradeoff relating construction cost to turnover rate, the predicted responses of optimal LMA to environment and size are flatter (Fig. 4). Given time, adaptation will fine tune the structure of each species, such they approach the biophysical limits of possibility within their average or typical environment (22) (Fig. 4a, black line). We assume biophysical limits reflect the original log-log-scaling tradeoff line. Indeed, empirical data supporting this trade-off are drawn from species average values (2). Now assume individuals attempting to construct leaves or wood with a trait value away from the trait value where selection has fine tuned that species suffer a penalty in the turnover rate that can be achieved (Fig. 4a), compared to what could be achieved via long-term evolution. Fig. 4 shows that the addition of such a penalty in the optimisation reduces the range of the optimal trait displayed by the species (Fig. 4, middle), bringing the observed trait response closer to what is empirically observed.

**Figure 4:**
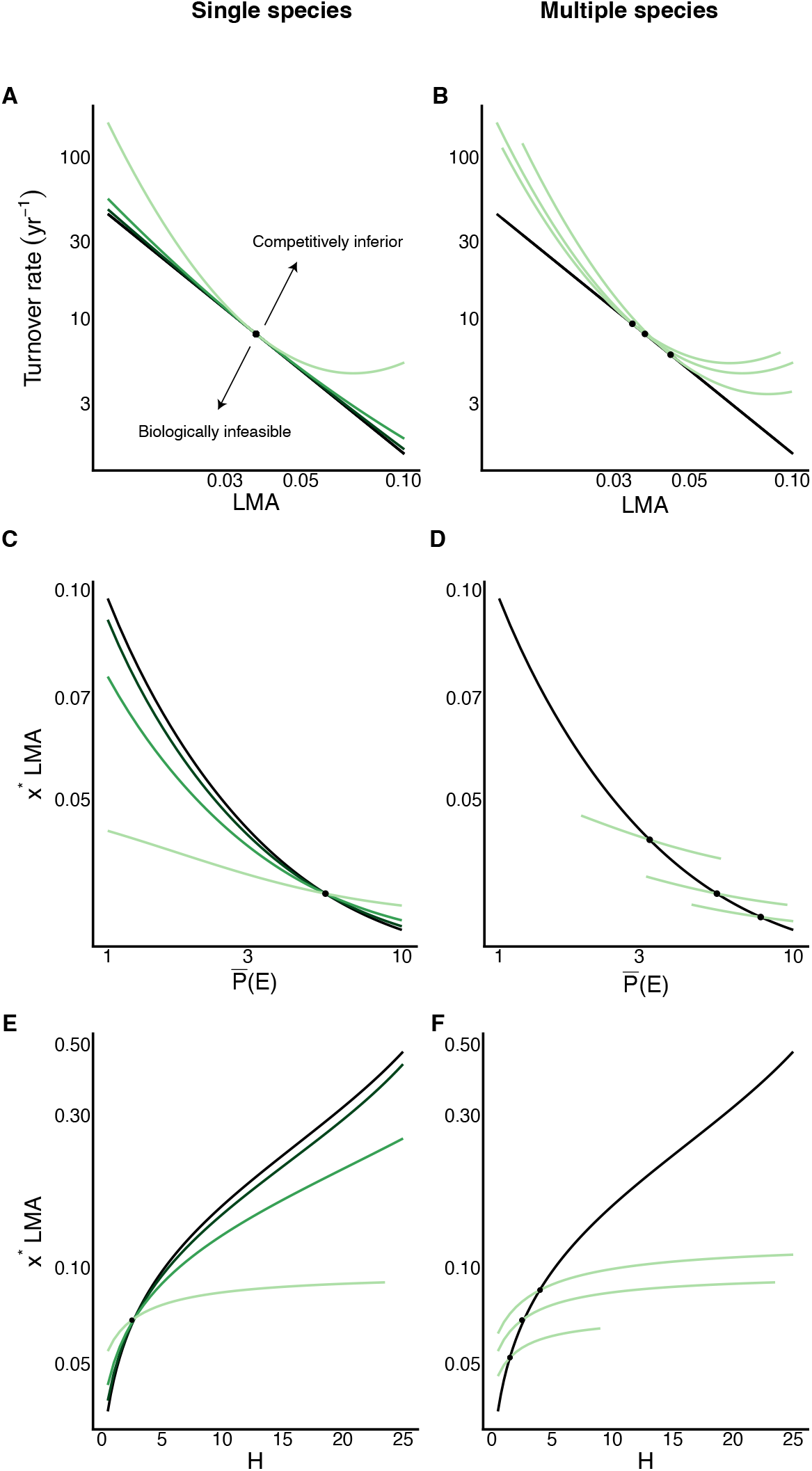
A modification of the basic model constrains the degree of plasticity within species along environmental gradients and through ontogeny. Left and right columns shows example for a single and multiple species, respectively. The top row shows the tradeoff associated with varying LMA, with the black line being the long-term biophysical limit, as documented in cross-species patterns in the leaf economics spectrum. Constructing leaves with trait combinations to the bottom-left of this line is hypothesised to be beneficial but biologically infeasible due to inevitable tradeoffs. Trait combinations to the top-right of this line are competitively inferior relative to the optimum value, all else being equal; being inefficient in either tissue construction costs, turnover rates or both. To this, we add a cost to plasticity (green lines), by applying a penalty on the turnover rate that can be achieved for a given LMA as a species attempts to construct tissues with a trait value away from it’s ideal value (dot). Lines show turnover with a small penalty (dark green) and strong penalty (light green) to plasticity. The right panel shows three species, each with a different optimal value of LMA, and a high penalty to plasticity. The second and third rows show the effect of plasticity costs on predicted traits with respect to changes in productivity (middle) and height (bottom). When penalties are introduced, the optimal trait achieved through a plastic response (green lines) diverges from the optimum that can be achieved by a species with either no penalty or perfectly adapted to a given location (black).

This constraint results in intra-specific variation in response to a change in productivity being more moderate than would otherwise be observed if such a species was instead replaced by another species which was optimally adapted to the new set of conditions (Fig. 4, right). This phenotypic extension to the basic model therefore provides a mechanistic explanation for how steep across-community responses can emerge from either or both species turnover or intra-specific responses curves, despite the latter exhibiting a flatter response (Fig. 4, right).

Past analyses based on TGT have demonstrated how the tradeoff between construction cost of leaves (LMA) and turnover leads naturally to coexistence of fast and slow-growing types, increases in LMA with reduced site productivity, how size moderates trait-based effects on growth, and how both traits respond to productivity or soil water (23; 10; 24). However, those findings relied on computationally expensive eco-evolutionary simulations. Here, we extend these findings by identifying a unifying underlying mathematical principle that explains these outcomes in a coherent mathematical framework.

Overall the proposed theory provides a much needed foundation for describing the effects of leaf, stem and root economics traits on plant performance and distribution. Although we did not explicitly consider root economic traits here, the framework can naturally be extended to other plant organs provided that it can be linked to a tradeoff between construction cost and photosynthesis or respiration rates in plant tissues. Our model does not capture all relevant processes shaping these traits, rather it illustrates the far-reaching consequences of a simple economics tradeoff for how evolutionarily expected trait values shift between sites and years, depending on other traits, and in response to size of individual.

## Methods

### Theory

The application of TGT here focusses on vegetative growth. An expanded version of equation 3 for investment in reproduction and the translation of leaf area into height can be found in (10). These additions are superfluous to the core results of the current paper and so were omitted.

Here, we make a single assumption about how traits impact function. We assume that for each tissue, making a unit area or volume of the tissue more cheaply reduces its longevity, eventually leading to higher turnover of that tissue. Thus, while a cheaper tissue increases the efficiency of leaf area deployment, it leads to a decrease in net biomass production. For leaves, this tradeoff has been empirically observed to follow a log-log scaling relationship between LMA and the rate of leaf turnover (*k*_l_) (2):

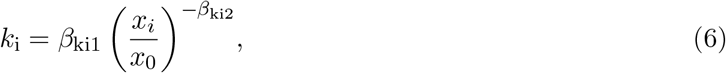

where *i ∈ l, s, r*, and (*β*_ki1_, *x*_0_, *β*_ki2_) are empirical constants. This tradeoff is the core of the well known leaf economics spectrum (2). We assume similar relationships exist for stem, and root tissues). The cost of turnover terms (*k*_*l*_, *k*_*s*_, *k*_*r*_) in turn influences the growth rate via their effects on an individual’s biomass production, given by:

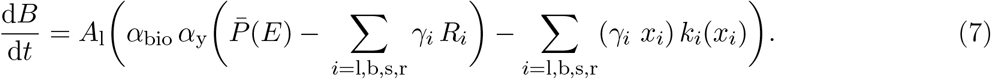

(See Table 2 for definitions). Eq. 7 states that net biomass production is given by the difference between photosynthetic income and costs, these being respiration and turnover of all plant tissues. We assume that respiration rates or each tissue are proportional to the area or volume of a tissue. As the functional components of a plant are related to its leaf area, respiration rates are therefore proportional to the area of leaf, the volume or area of each tissue per unit leaf area (*γ*_*i*_) and the tissue specific rates (*R*_*i*_) per unit area or volume, which are not influenced by economics traits. Turnover, by contrast, is influenced by the economics traits. Those economics traits translate areas and volumes into mass, and also influence the turnover rates *k*_*i*_.

Solving for an optimum requires solving for the value of *x*^∗^ where Eq. 5 is satisfied. This point was obtained via bisection search.

Note that eq. 5 also captures situations where a trait only impacts 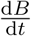 and has no impact on the efficiency of leaf area deployment. In this case, both terms on the LHS of Eq. 5 are zero. The only way to satisfy the optimum criteria is when 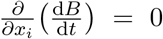. This criterion is the same as would be obtained if the optimum were solved directly with respect to biomass production, rather than growth rate. Hence, optimising biomass production is a special case of the more general result optimising leaf area growth rate.

For Figs 1-2 the model was run assuming plants were able to adjust traits with no penalty, i.e. so that turnover followed Eq. 6. The model was extended to consider the effects of plasticity in traits, by applying a penalty to the turnover that could be realised by the plant, with the magnitude of the penalty based on the divergence of the trait from an ideal value, where investment is most effective. To implement this modification, we assumed each species had an ideal trait value *x*_*c*_. Turnover was then given by

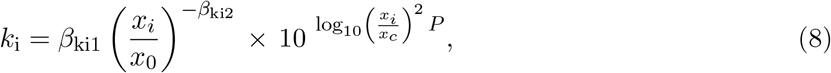

where *P* governs the rate *k* diverges from the value given in Eq. 6, as *x* moves away from *x*_*c*_. Note that when *x* = *x*_*c*_ Eq. 8 collapses to give Eq. 6. Importantly, sub-optimum turnover rates are just one mechanism which could hypothetically limit the plasticity of traits, and there are likely many biophysical mechanisms. Consequently, the central aim of the penalty was to capture as an emergent property the phenomenological observation that intra-specific trait variation limits a given taxon’s ability to persist along an environmental gradient beyond a certain point, leading to species turnover.

### Implementation

The model was implemented as the FF16 physiological module within Version 2.0 of the ‘PLANT’ package (25) for R, using additional parameters for the relationship between height and leaf area from (10). The ‘PLANT’ package makes use of supporting packages ‘RCPP’ (26) and the Boost Library for C++ (27). All analyses were conducted in R ver 4.5.0.

## Abbreviations

LMA: Leaf mass per unit leaf area
SSD: Specific stem density
TGT: Trait growth theory
EEO: Eco-evolutionary optimality
LHS: Left-hand side
RHS: Right-hand side

## Acknowledgements

We thank I Wright, F Kar and T O’Brien for useful discussions. IT was supported by grants from the Australian Research Council (DP200100555, LP230100049).

## References

[1] Westoby M, Falster DS, Moles AT, Vesk P, Wright IJ (2002) Plant ecological strategies: Some leading dimensions of variation between species. Annual Review of Ecology and Systematics 33:125–159.

[2] Wright IJ, et al. (2004) The worldwide leaf economics spectrum. Nature 428(6985):821–827.

[3] Chave J, et al. (2009) Towards a worldwide wood economics spectrum. Ecology Letters 12(4):351–366.

[4] Díaz S, et al. (2016) The global spectrum of plant form and function. Nature 529:167–171.

[5] Franklin O, et al. (2020) Organizing principles for vegetation dynamics. Nature Plants 6(5):444–453.

[6] Westoby M, Cornwell WK, Falster DS (2012) An evolutionary attractor model for sapwood cross section in relation to leaf area. Journal of Theoretical Biology 303:98–109.

[7] Wolf A, Anderegg WRL, Pacala SW (2016) Optimal stomatal behavior with competition for water and risk of hydraulic impairment. Proceedings of the National Academy of Sciences p. 201615144.

[8] Wang H, et al. (2023) Leaf economics fundamentals explained by optimality principles. Science Advances 9(3):eadd5667.

[9] Falster DS, Brännström Å, Dieckmann U, Westoby M (2011) Influence of four major plant traits on average height, leaf-area cover, net primary productivity, and biomass density in single-species forests: a theoretical investigation. Journal of Ecology 99(1):148–164.

[10] Falster DS, Duursma RA, FitzJohn RG (2018) How functional traits influence plant growth and shade tolerance across the life cycle. Proceedings of the National Academy of Sciences 115(29):E6789–E6798.

[11] Fisher RA, et al. (2018) Vegetation demographics in Earth System Models: A review of progress and priorities. Global Change Biology 24(1):35–54.

[12] Duursma RA, Falster DS (2016) Leaf mass per area, not total leaf area, drives differences in above-ground biomass distribution among woody plant functional types. New Phytologist 212(2):368–376.

[13] Towers IR, et al. (2024) Revisiting the role of mean annual precipitation in shaping functional trait distributions at a continental scale. New Phytologist 241(5):1900–1909.

[14] Reich PB (2014) The world-wide ‘fast–slow’ plant economics spectrum: A traits manifesto. Journal of Ecology 102(2):275–301.

[15] Westoby M, Schrader J, Falster D (2022) Trait ecology of startup plants. New Phytologist 235(3):842–847.

[16] Fernández-de-Uña L, Martínez-Vilalta J, Poyatos R, Mencuccini M, McDowell NG (2023) The role of height-driven constraints and compensations on tree vulnerability to drought. New Phytologist 239(6):2083–2098.

[17] Hietz P, Valencia R, Joseph Wright S (2013) Strong radial variation in wood density follows a uniform pattern in two neotropical rain forests. Functional Ecology 27(3):684–692.

[18] Fischer FJ, et al. (2025) Beyond species means – the intraspecific contribution to global wood density variation. bioRxiv.

[19] Poorter L, Bongers F (2006) Leaf Traits Are Good Predictors of Plant Performance Across 53 Rain Forest Species. Ecology 87(7):1733–1743.

[20] Reich PB, Ellsworth DS, Uhl C (1995) Leaf carbon and nutrient assimilation and conservation in species of differing successional status in an oligotrophic Amazonian forest. Functional Ecology 9(1):65–76.

[21] Auld JR, Agrawal AA, Relyea RA (2010) Re-evaluating the costs and limits of adaptive phenotypic plasticity. Proceedings of the Royal Society B: Biological Sciences 277(1681):503–511.

[22] Reich PB, et al. (2003) The evolution of plant functional variation: Traits, spectra and strategies. International Journal of Plant Sciences 164(S3):S143–S164.

[23] Falster DS, Brännström Å, Westoby M, Dieckmann U (2017) Multitrait successional forest dynamics enable diverse competitive coexistence. Proceedings of the National Academy of Sciences 114(13):E2719–E2728.

[24] Towers IR, O’Reilly-Nugent A, Sabot MEB, Vesk PA, Falster DS (2024) Optimising height-growth predicts trait responses to water availability and other environmental drivers. Plant, Cell & Environment 47(12):4849–4869.

[25] Falster DS, FitzJohn RG, Brånnström A, Dieckmann U, Westoby M (2016) plant: A package for modelling forest trait ecology and evolution. Methods in Ecology and Evolution 7(2):136–146.

[26] Eddelbuettel D, Francois R (2011) Rcpp: Seamless R and C++ Integration. Journal of Statistical Software 40(8).

[27] Schäling B (2014) The Boost C++ Libraries. (XML Press, Laguna Hills, Calif), 2nd english ed edition.

[28] Falster DS, Westoby M (2005) Alternative height strategies among 45 dicot rain forest species from tropical Queensland, Australia. Journal of Ecology 93(3):521–535.

[29] Stahl U, et al. (2013) Whole-plant trait spectra of North American woody plant species reflect fundamental ecological strategies. Ecosphere 4(10):art128.

[30] Domínguez MT, et al. (2012) Relationships between leaf morphological traits, nutrient concentrations and isotopic signatures for Mediterranean woody plant species and communities. Plant and Soil 357(1):407–424.

[31] He D, Chen Y, Zhao K, Cornelissen JHC, Chu C (2018) Intra- and interspecific trait variations reveal functional relationships between specific leaf area and soil niche within a subtropical forest. Annals of Botany 121(6):1173–1182.

[32] ter Steege H, et al. (2025) Functional composition of the Amazonian tree flora and forests. Communications Biology 8(1):355.

[33] Umaña MN, et al. (2021) Shifts in taxonomic and functional composition of trees along rainfall and phosphorus gradients in central Panama. Journal of Ecology 109(1):51–61.

[34] Cornwell WK, Ackerly DD (2009) Community assembly and shifts in plant trait distributions across an environmental gradient in coastal California. Ecological Monographs 79(1):109–126.

